# Constraints on avian seed dispersal reduce potential for resilience in degraded tropical forests

**DOI:** 10.1101/2023.09.09.556978

**Authors:** Jack H. Hatfield, Cristina Banks-Leite, Jos Barlow, Alexander C. Lees, Joseph A. Tobias

## Abstract

1. Seed dispersal – one of the many services supplied by biodiversity – is a critical process underpinning the resilience of tropical forests. Forest loss or degradation typically leads to defaunation, altering seed transfer dynamics and impairing the ability of forested habitats to regenerate or recover from perturbation. However, the extent of defaunation, and its likely impacts on the seed dispersers needed to restore highly degraded or clear-felled areas, remains poorly understood, particularly in human-modified tropical forest landscapes.

2. To quantify defaunation of seed-dispersing birds, we used field survey data from more than 400 transects in three regions of Brazil, first comparing the recorded assemblages with those predicted by geographic range maps, and then assessing frugivore habitat associations across gradients of land cover modification at local scales.

3. We found that current bird assemblages have lower functional trait diversity than predicted by species range maps in Amazonia (4–6%), with a greater reduction (28%) for the Atlantic Forest region, which has been more heavily deforested for a longer period. These reductions are probably caused by local extinctions of forest-dependent bird species following land-use change.

4. Direct measures of seed dispersal are difficult to obtain, so we instead focused on the potential for seed transfer inferred from shared species occurrence between land cover types. Of 83 predominantly frugivorous bird species recorded in relatively intact forests, we show that 10% were absent from degraded forest, and 57% absent from the surrounding matrix of agricultural land covers, including many of the large-beaked species. Of 112 frugivorous species using degraded forest, 47% were absent from matrix habitats.

5. Our findings suggest that degraded forest can supply seed dispersal services to adjacent cleared lands, and that direct transfer of seeds from intact forest to cleared areas may be limited, particularly for large-seeded trees. We conclude that resilience of tropical forest landscapes is best achieved by protecting a mosaic of forest types, including sufficient core areas of intact forest surrounded by buffer zones of degraded forest.

## 1 INTRODUCTION

The degradation and fragmentation of tropical forests, along with their conversion to production landscapes, is driving widespread defaunation (Canale et al., 2012; Dirzo et al., 2014) with large frugivorous birds particularly affected (Bovo et al., 2018; Bregman et al., 2014). This loss of species can disrupt seed-dispersal networks (da Silva & Tabarelli, 2000), thereby limiting forest regeneration (Galetti et al., 2013; Gardner et al., 2019). Defaunation alters assemblages of frugivores in disturbed forests, with few species occurring in adjacent non-forest landscapes (Bregman et al., 2016). This potentially impedes recolonisation by tree species that have become extirpated, reducing ecosystem resilience and raising the cost of reforestation programmes (Chazdon & Uriarte, 2016). Few studies, however, have assessed how habitat degradation affects potential seed transfer from intact tropical forests to adjacent cleared areas by examining frugivore communities.

One possibility is that many seed-dispersers associated with intact forests rarely if ever visit non-forest habitats, such that species transferring seeds into matrix habitats are primarily those associated with degraded forests and forest edges. If this is the case, tree species recruitment will be adversely affected because these frugivorous birds are a non-random subset in terms of key ecological traits, such as beak or gape size (Bovo et al., 2018). In particular, the absence of species with larger beaks and wider gapes may constrain seed transfer because the ability of avian frugivores to disperse seeds is limited by gape width, which places an upper physiological constraint on the size of seeds they can ingest whole and hence which tree species can be dispersed (Burns, 2013; Wheelwright, 1985). This trait-matching between avian frugivores and their food plants offers a useful tool for assessing the structure and function of mutualistic interaction networks (Dehling et al., 2014; McFadden et al., 2022) and their response to environmental change (Schleuning et al., 2020).

In accordance with the trait-matching hypothesis, the local extinction of large-gaped birds has been shown to impair dispersal of larger seeds (da Silva & Tabarelli, 2000; Galetti et al., 2013). These local extinctions of key seed dispersers may lead to declining plant populations, particularly for large-seeded tree species, resulting in changes to recruitment and composition (Sethi & Howe, 2009). Movement constraints on seed-dispersers may also cause the rewiring of interaction networks between seed dispersers and their food plants, with unpredictable outcomes driven by species-specific responses to different land covers (Habel et al., 2019; Rehm et al., 2017). In effect, the type of seeds dispersed between different land cover types will be governed by the traits of frugivores that can persist in or routinely disperse through both forest and non-forest landscapes. In addition, this process of seed transfer may involve an extra step from intact to degraded forest and then to matrix simply because intact forests are usually separated from the matrix by an intervening buffer of degraded or secondary forests (Mayhew et al., 2019; Nunes et al., 2022).

To determine the effects of disturbance on seed dispersal, two key questions must be answered. First, given that the potential to disperse different seeds is dependent on species traits, it is important to examine the functional diversity (FD) of the bird species assemblage. FD metrics use traits to quantify the potential roles played by species in a given community (Cadotte et al., 2011; Petchey & Gaston, 2002). One way to examine FD is with a volume-based measure (Mason et al., 2005) with morphological traits providing a representation of ecological niche space (Pigot et al., 2020; Tobias et al. 2020). If functionally unique species are lost, the functional volume is reduced, indicating the likely impairment of seed dispersal services for certain plant species. If, however, the species lost are not functionally unique, this can still result in reduced functional redundancy, i.e., where different species perform similar roles, species loss does not necessarily reduce FD but nonetheless leads to an increase in future risk by leaving seed dispersal service reliant on fewer species (Biggs et al., 2020). Second, it is important to understand how frugivores use the modified landscape. If certain species or traits are restricted to a single land cover type, seed dispersal between different land covers will also be restricted, as birds are not able to act as “mobile links’’ (Sekercioglu, 2006), which will in turn reduce the potential for forest regeneration.

In this study, we aim to investigate how land-use change and habitat degradation affects the potential for natural seed dispersal by assessing changes in key traits and FD in assemblages of fruit-eating birds. To investigate the potential functional impacts of defaunation at a regional scale, we compare the species assemblages predicted by geographical range maps to those recorded by intensive field surveys using the distribution of individual traits and measures of functional volumes (which account for trait combinations). Previous research shows local declines of larger-beaked seed-dispersers in tropical forests (Galetti et al. 2013; Dirzo et al., 2014; Pérez-Méndez et al., 2016; Bovo et al. 2018) because of a range of factors, including slower reproductive output, larger spatial requirements and sensitivity to hunting. We therefore predict that assemblages recorded by recent surveys will have fewer large species than predicted by range maps, contributing to a reduction in FD.

To examine different seed-dispersal scenarios, we subdivide land cover into three broad classes – intact forest, degraded forest and the agricultural/silvicultural matrix. Previous work has identified high levels of species turnover across land cover types in the same assemblages (Hatfield et al., 2020; Moura et al., 2013; Solar et al., 2015) and elsewhere (González-Varo et al., 2017), suggesting that direct transfer of seeds between very different habitats is reduced. Because of this and the pattern of extirpation of larger species we predict that movements between intact forest and the matrix are potentially limited to a reduced shared species pool representing a narrower range of traits, whereas there is a higher potential for seed dispersal between intact and degraded forests, or between degraded forests and matrix (Figure 1).

**FIGURE 1.**
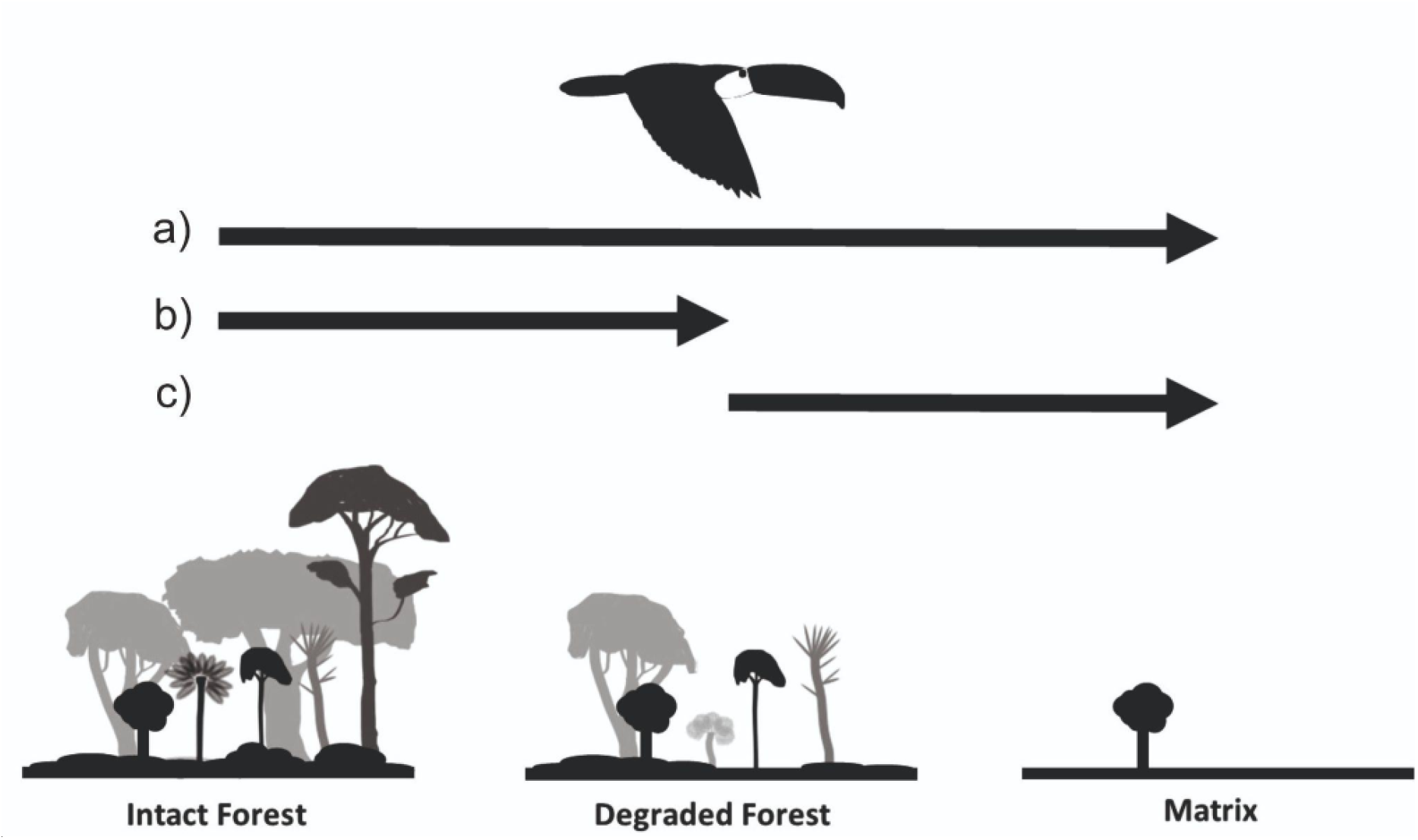
Schematic of possible seed dispersal scenarios from intact forest to matrix. Seeds could be transferred directly from intact forest into the matrix (Scenario a); transferred only from intact forest to degraded forest without subsequent seed dispersal into the matrix (Scenario b); or only transferred from degraded forest to the matrix without any prior seed dispersal from intact to degraded forest (Scenario c). Alternatively, seeds can be transferred from intact forest to surrounding matrix via a stepwise pathway (b then c), although this involves a major rate-limiting step – that is, it can take many years before seeds deposited in degraded forest develop into trees mature enough to bear fruit that can be consumed and subsequently transferred into the matrix.

## 2 MATERIALS AND METHODS

### 2.1 Bird surveys and comparison with historical avifauna

We analysed avifaunal data collected from the state of São Paulo (Brazilian Atlantic Forest; 23°S 45°W) and two areas in the state of Pará (Brazilian Amazonia): Santarém (03°S 55°W) and Paragominas (03°S 48°W). These three survey sites are each in a different biogeographic province and differ in their disturbance histories, providing a useful comparison. All datasets used three 15-minute point counts spaced along a transect to sample bird assemblages but differed slightly in overall study design (owing to different habitat extent and configuration). Amazonian sampling was conducted in 2010–2011 at 352 transects (300 m, 75 m radius points 150 m apart, surveyed twice; Supplementary Methods; Lees et al., 2012, 2013). Atlantic Forest sampling was conducted in 2015 –2017 at 147 transects (150 m, 25 m radius points 75 m apart, surveyed four times; Supplementary Methods; Hatfield et al., 2020).

All study regions have undergone dramatic forest loss and fragmentation (Figure S1; Gardner et al., 2013; Joly et al., 2014), leading to the local extinction of bird species, and even – in the case of the Atlantic Forest – global extinctions (Brooks et al., 1999; Lees & Pimm, 2015). To compare current bird assemblages with that expected in each region prior to large scale impacts, we compiled a list of species historically present according to published range maps for each region (BirdLife International and Handbook of the Birds of the World, 2020). We determined which native, introduced and (locally) extinct bird species had geographical distributions overlapping the sampling sites, independently for the three regions. Thus, providing a full potential species list for each after taxonomic alignment and exclusions based on finer scale biogeographic factors (Supplementary Methods).

### 2.2. Trait data and ecological classifications

We restricted our sample of study species to those with diets composed ≥ 50% of fruit according to a previous scoring system (Wilman et al., 2014). By omitting species with a smaller proportion of fruit in their diet we may exclude some generalist taxa that play important roles in dispersing seeds into matrix areas. However, our approach means that we focus primarily on obligate and near obligate frugivores which form the core of seed dispersal networks (de Assis Bomfim et al., 2018), due to their higher consumption and visitation rates (Pigot et al., 2016). We did not exclude species that are often classed as seed predators (e.g., parrots), as those with a large percentage of fruit in their diet (≥ 50%) have been shown to provide some seed dispersal services (Heleno et al., 2011; Tella et al., 2015).

Following disturbance and clearance of forests, the likelihood that a species is lost from an assemblage is related to its forest dependency. We therefore classified species as being forest and non-forest associated for the FD analyses, with all species initially analysed, followed by the forest species subset alone. Classification used the primary habitat category in AVONET (Tobias et al., 2022); in addition, we re-ran the analyses based on an independent classification, focusing on species listed as having medium or high forest dependency by BirdLife International (BirdLife International, 2022; Buchanan et al., 2011) with clear discrepancies reviewed (Supplementary Methods).

To calculate FD, we used a set of morphological traits (Table S3) associated with diet (beak length, width and depth), locomotion (tarsus and tail length) and dispersal (wing length and hand-wing index). Species averages for these traits were based on published measurements of museum specimens and live-caught birds (Sheard et al., 2020; Tobias et al., 2022; mean sample per species = 16.6 individuals). Hand-wing index (HWI) is a metric of wing shape associated with flight efficiency and dispersal distance (Sheard et al. 2020). We also added average gape size for all species (McFadden et al. 2022; mean sample per species = 17.7 individuals). For eight species not included, we used previously unpublished gape size values measured following the same methodology (Supplementary Methods).

### 2.3 Assemblage functional diversity

We estimated FD using the convex hull volume of the functional space occupied by an assemblage (Villéger et al., 2008). The functional space was constructed in three dimensions using axes based on the total range of traits across all avian frugivores in each region independently (combining the species lists from transects and geographical range maps).

Functional axes were generated using a two-step PCA process designed to partition the effects of body size from morphological trait variation associated with foraging and locomotion (Trisos et al., 2014). Specifically, two PCAs were initially conducted – a PCA based on foraging traits (gape size, beak length along the culmen, beak length to the anterior edge of the nares, beak width and beak depth) and a second based on locomotion traits (tarsus length, tail length, wing length and HWI). The first axis from both the foraging and locomotion trait PCAs were then combined in a second PCA that reflects variation in body size. This provided the three axes to construct the functional space, one from the second step PCA reflecting variation in body size, the second axis from the foraging PCA reflecting variation in trophic niche and the second axis from the locomotion PCA reflecting variation in locomotion (Figure S2). FD calculations were conducted in the R package ‘betapart’ (Baselga et al., 2021).

This analysis of FD allowed us to compare species assemblages predicted by range maps and those recorded by surveys. The list derived from range maps provides an approximation of the expected species assemblage. The proportion of the functional volume of the expected species assemblage (inferred from range maps) that was retained (shared) and lost by the observed assemblage (inferred from site transects) was calculated. The percentage retained was then also compared to that retained by randomly generated assemblages (with richness held constant; Supplementary Methods). This allowed us to examine whether the species losses estimated from the survey data reduced FD retention to a greater extent than a trait-independent process of loss whereby species are lost at random with respect to their traits as opposed to a trait-based filter (e.g., loss of the largest species). All estimates were conducted for the full avian frugivore assemblage and then for a restricted sample of forest species.

### 2.4 Assessing the potential for seed transfer among land cover types

When assessing the potential for seed transfer among land cover types we chose to focus primarily on frugivore gape width as this provides the best measure of seed dispersal limitation with regards to size. The land cover classifications provided in the original studies (Hatfield et al., 2020; Moura et al., 2013) were aggregated into three categories: intact forest, degraded forest and matrix (Table S1 and S2). Intact forest is used as a relative term as these landscapes are still likely affected by historic and ongoing disturbance.

First, we compiled frugivore lists for each land cover type (each region independently). Next, we produced lists of species common to both intact forest and degraded forest then, intact forest and the matrix. We also produced a list of species common to both degraded forest and the matrix. This allowed us to calculate the percentages of species in common and to evaluate possible seed dispersal pathways in the context of gape width, with the overlap lists corresponding to the potential steps in Figure 1. Species overlaps were also calculated when sampling effort across land cover types was fixed (Supplementary Methods). To include estimates of dispersal ability, we also repeated the overlap comparison with HWI and with body mass (AVONET; as an indication of body size). The proportion of functional diversity not retained by the species shared between land cover types was also estimated by applying the above methods (2.3) to lists of species occurring in multiple habitat combinations (Supplementary Methods). For both gape width and FD, we compared the observed species overlaps to random selections to test for trait-based filters (such as an increased likelihood of extirpation for species with large body size; Supplementary Methods).

In addition to the percentage overlaps for each region, overall figures were calculated by combining the species by land cover lists from the three regions, then calculating overlaps. This approach differs from averaging values across regions as some species are found in multiple regions and each region has a different species richness. Land cover associations may also differ between regions as sensitivity to forest degradation is not uniform across a species’ geographic range (Moura et al., 2016; Orme et al., 2019). This means that across-region pooling may not accurately reflect the associations of individual species in a particular region. On the other hand, many of these species have few detections even with the large sampling effort considered, so the pooled figures represent an increased sampling effort for species found in multiple regions.

## 3 RESULTS

### 3.1 Functional diversity of frugivore assemblages

Field surveys of frugivorous birds detected 73 species in Santarém, compared with 104 predicted by geographical range maps for the region. Similarly, surveys detected 76 species compared to 87 species predicted in Paragominas, and 46 species compared to 81 species predicted in São Paulo. The species assemblages recorded by field surveys in both Amazonian study regions suggested a loss of ≤ 6% of FD compared to original levels of FD calculated for the same regions based on range maps. The equivalent reduction in FD for São Paulo was much higher, with an estimated decline of 28% compared with expected FD calculated from range maps. Similar results were found when the assemblage was limited to only forest species (Table 1 and Figures 2, S3-4). When comparing observed assemblages to those randomly generated from the species pool, we found no strong evidence of losses being clustered in relation to traits (Table 1).

**FIGURE 2.**
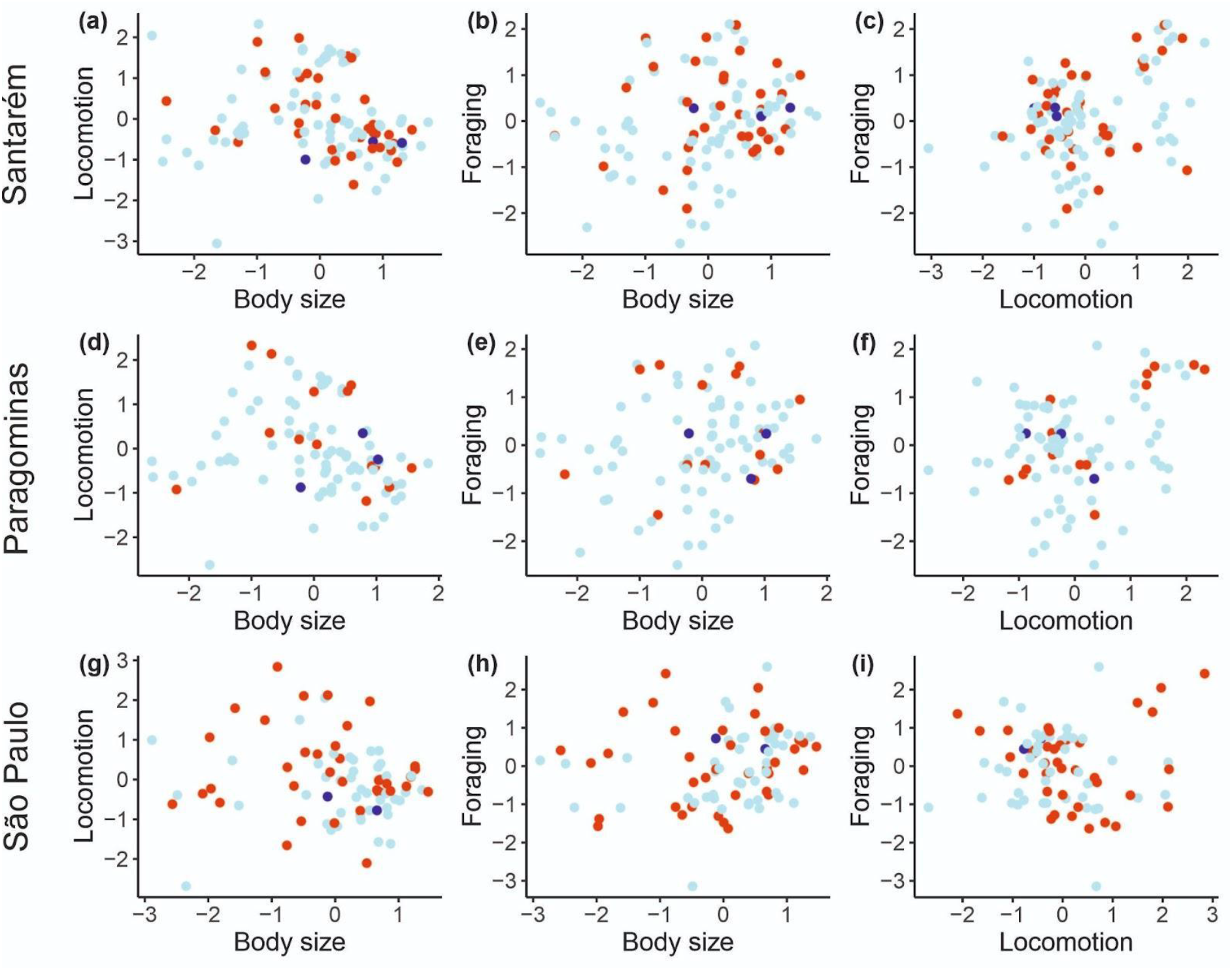
Distribution of species along trait axes for all frugivores in Santarém (a, b and c), Paragominas (d, e and f) and São Paulo (g, h and i). Data points are coloured to indicate whether species in the frugivore assemblages are predicted to occur based on geographic distributions but not detected by surveys (red), found in surveys but not predicted to occur by geographic range distributions (dark blue), or both detected by empirical surveys and predicted by geographic distribution (pale blue).

**TABLE 1:** Historical declines in functional diversity (FD) for study assemblages. Declines were estimated as a percentage of the expected FD calculated from the projected historical avifauna determined by geographical range maps. Values are given for each of the three regions (Santarém, Paragominas and São Paulo) for all frugivores and for forest frugivores (i.e., frugivores largely restricted to forest vegetation). Results for forest frugivores are shown for two different classifications: main values are based on species with primary habitat classified as forest in AVONET (Tobias et al. 2022); values in brackets are based on BirdLife International’s forest dependency categories (BirdLife International, 2022; Buchanan et al., 2011).

| Region | All frugivores |  | Forest frugivores |  |
| --- | --- | --- | --- | --- |
| | Percentage<br>FD reduction | Randomly<br>generated FD<br>reduction ≤<br>observed loss (%) | Percentage<br>FD reduction | Randomly<br>generated FD<br>reduction ≤<br>observed (%) |
| Santarém | 6 | 6 | 7 (8) | 12 (17) |
| Paragominas | 4 | 35 | 2 (1) | 19 (22) |
| São Paulo | 28 | 32 | 33 (33) | 48 (29) |

Considering individual traits (Table S4–S6; Figure S5–S7), the Santarém survey assemblage had lower maximum values for beak length and HWI compared to that predicted from range maps. The mean (± sd) gape size was similar for the field survey (15 mm ± 7.8 mm) and range map (14.7 mm ± 7.7 mm) assemblages. For Paragominas, only HWI differed in maximum values, with a lower maximum in the field survey assemblage. The mean gape size was similar for the field survey (14.7 mm ± 7.0 mm) and range map (14.5 mm ± 6.8 mm) assemblages. In São Paulo, observed maximum values for tarsus length, wing length and HWI were lower, and most traits showed reduced maximum values for the forest species assemblage comparisons. For this region there was a slight reduction in the mean gape size when comparing the field survey (11.6 mm ± 6.8 mm) to the range map (12.5 mm ± 6.9 mm) assemblage.

### 3.2 Potential for seed dispersal

Across study regions, we found that 57% of species found in intact forest were not detected in the matrix, and are therefore highly unlikely to disperse seeds between the two (Figure 1a). Regionally the number of species found in intact forest but not in the matrix was 28 (62% of intact forest assemblage) in Santarém, 30 (79%) in Paragominas and 12 (46%) in São Paulo. In all three regions, only a few species present in intact forest were absent from nearby degraded forest (Figure 1b; 10% of frugivore species richness in intact forest across all sites): 1 species (2%) in Santarém; 4 species (11%) in Paragominas; 6 species (23%) in São Paulo. Roughly half (47% overall) of frugivore species were found in degraded forest but not in the nearby matrix (Figure 1c): 38 species (54% of the degraded forest assemblage) in Santarém; 48 species (70%) in Paragominas; 12 species (32%) in São Paulo.

The FD represented by the species shared between land covers did not correspond exactly with species richness. The species shared between intact forest and matrix did not retain 59%, 90% and 60% of the intact forest FD in Santarém, Paragominas and São Paulo, respectively (Table S7). The high species overlap between intact and degraded forest translated into lower FD losses with 16%, 28% and 1% (Santarém, Paragominas and São Paulo; Table S7) of intact forest assemblage FD not retained by the species shared between intact and degraded forest. When considering the species shared between degraded forest and matrix, 52%, 62% and 33% (Santarém, Paragominas and São Paulo; Table S7) of degraded forest assemblage FD was not retained. When considering specific traits, very few large-gaped frugivore species occurred across the full land cover gradient (Figure 3), with similar patterns observed for HWI and body mass (Figures S8 and S9).

**FIGURE 3.**
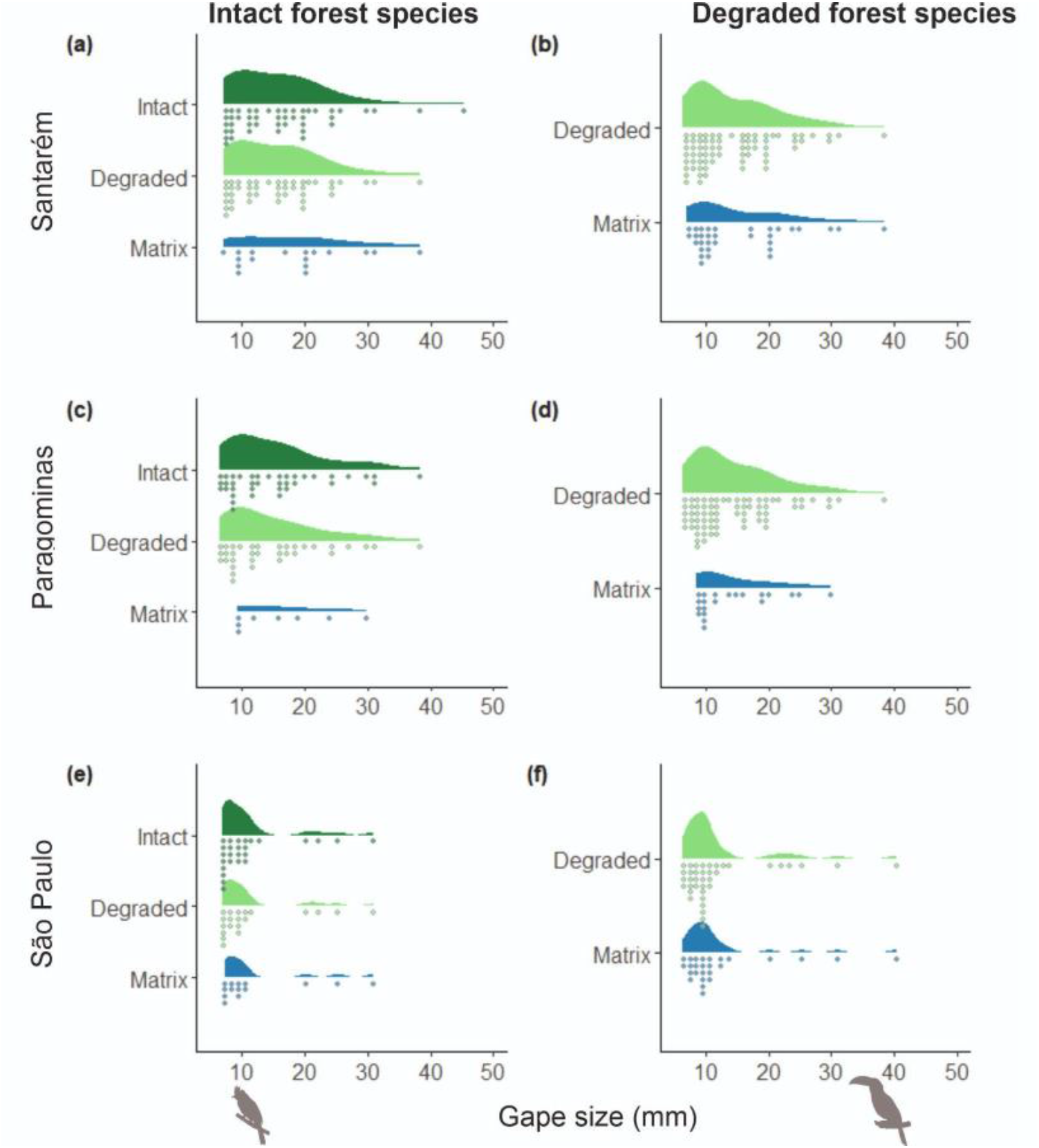
Gape sizes of frugivorous birds across different land cover types for Santarém (a, b), Paragominas (c, d) and São Paulo (e, f). Gape size (a horizontal linear measurement across the base of the mandibles where they join) is an index of maximum fruit size that can be consumed by each species. Each dot represents a single species, grouped into 1 mm bins. Left-hand panels (a, c, e) show species recorded in intact forest, and indicate whether they also occur in degraded and matrix habitats; right-hand panels (b, d, f) show species recorded in degraded forest, and indicate whether they also occur in the matrix. Bird species with larger gape sizes (>20 mm) are most frequent in intact forest and rarer in degraded or matrix habitats. Silhouettes of a small-gaped *Myiarchus* flycatcher and a large-gaped toucan (Ramphastidae) were obtained from phylopic.org.

Given that the intensity of surveys differed across land cover types (Table S2), comparisons among bird assemblages may be influenced by sampling biases. However, when we ran the analyses with sampling effort fixed across land cover types, we found that overlap percentages were qualitatively similar (Figure S10). In addition, given that in many cases few species were shared between land cover types, we tested whether any reductions in maximum gape width and FD could be attributed to trait-based filtering or were indistinguishable from random nestedness. Evidence for trait-based filtering was only found when considering FD in both Amazonian regions (Table S7) and also for the gape width of species shared between intact and degraded forest in Santarém (Table S8).

## 4 DISCUSSION

Our results suggest that seed dispersal between intact tropical forest and matrix landscapes will proceed through a combination of direct and stepwise processes (scenarios a–c; Figure 1). Given the high species turnover observed across land cover gradients, we conclude that relatively few frugivore species are involved in direct seed dispersal from remaining patches of intact forest into the surrounding matrix, with these events likely to be rare. Moreover, we can infer that direct dispersal is mainly restricted to smaller seeds, because a relatively low number of large-gaped frugivorous bird species are shared between both habitats. This finding implies that large-seeded tree species are more likely to require a stepwise process, potentially delaying natural forest regeneration by decades if the distance to intact forest is too large.

The number of avian frugivores shared between more structurally similar habitats (that is, shared between intact and degraded forests; or between degraded forests and the matrix) is relatively high. Additional seed dispersal between intact forest and matrix may therefore follow a stepwise pathway through degraded intermediaries, such as secondary forests. This stepwise dispersal pattern is much less efficient, however, as it requires seeds deposited in intermediate habitats to develop into seed-bearing trees before they can be transferred to the surrounding matrix. These results also indicate that transport of larger seeds by birds is reliant on a relatively small number of large-gaped species (Burns, 2013; Wheelwright, 1985), providing an important insight into likely challenges for natural regeneration in human-modified tropical forest landscapes.

Reliance on few large-gaped species as potential dispersal vectors between intact forest and matrix is clearest for the sites in São Paulo and Paragominas but can still be seen to a lesser extent in Santarém (Figure 3). In São Paulo and Paragominas, it is not only that there are few large-gaped species shared between intact forest and matrix, very few large-gaped species are recorded in the matrix at all (Figure S11). These findings align with previous studies reporting that relatively few bird species use both intact and cleared land-cover types (Rehm et al., 2017) and that these ecologically flexible species with the greatest potential to act as mobile links for seed dispersal tend to be smaller bodied with smaller gapes (Pizo & dos Santos, 2011).

In our Amazonian landscapes, the dispersal of very large seeds between intact forest and the cleared matrix relies on species like *Pteroglossus aracari* and other toucans (*Ramphastidae*). In the Atlantic Forest landscapes, birds with larger gape sizes detected in both intact forest and cleared matrix included toucans (*Ramphastos dicolorus*) and cotingas (*Pyroderus scutatus*). While the importance of large-gaped frugivorous birds for the recruitment of large seeds is well known (Beltrán & Howe, 2020; Reid et al., 2021), reliance on very few species reduces resilience to future perturbations, particularly if population size is also low. Our results show that, especially for São Paulo, most large-gaped species are either rare visitors to matrix areas or missing from these landscapes altogether.

All our surveys detected fewer bird species than predicted by range maps, resulting in slightly lower estimates of FD than anticipated from geographical ranges. In contrast with some previous studies (e.g., Bovo et al, 2018), we found no evidence that larger-gaped species were more likely to drop out from current assemblages. Nonetheless, their loss has a disproportionate impact on FD because redundancy declines towards the extremes of the trait distribution. The absence of one large-gaped species therefore represents a greater decline in ecological function and resilience than the loss of a different species more centrally placed on the gape-size gradient, where redundancy is high (Ali et al. 2023).

### 4.1 Methodological limitations

It is worth emphasising that distribution maps can provide misleading information about historical assemblages. Mapping exercises tend to overestimate species richness at finer scales (Hurlbert & Jetz, 2007), and are unlikely to account for habitat specialisation (Jetz et al., 2008). In addition, field surveys are subject to imperfect detection of species with cryptic behaviours or spatiotemporally patchy distributions. Even with intensive sampling effort, the diversity of rare or inconspicuous species in tropical bird communities makes detection challenging (Banks-Leite et al., 2014; Robinson et al., 2018). Nonetheless, the number of species overlooked by our field surveys is likely to be very low, particularly for large-gaped species which tend to be highly conspicuous when present. Almost all the losses we detect are therefore likely to represent true losses of function or declines in functional redundancy (Figure 2). In addition, even the frugivorous species detected in our study landscapes may be greatly reduced in population size by habitat loss and hunting, meaning that the volume and distance of seed dispersal may be substantially reduced in comparison with historical landscapes.

Our analyses focus exclusively on specialist avian frugivores, potentially overlooking the contribution of bird species for which fruit is not a major part of their diet, such as omnivores (Carlo & Morales, 2016; Rehm et al., 2017) or invertivores that supplement their diet with some fruit (Camargo et al., 2020). Migratory tyrant-flycatchers, for instance, may consume large quantities of berries in degraded Amazonian and Atlantic forests during the non-breeding season, with potentially large impact on early successional stages (Athiê & Dias 2016). Other taxa such as fish, reptiles and mammals can disperse seeds but again may perform different roles in the dispersal system (de la Peña-Domene et al., 2014; Donatti et al., 2011). Moreover, they face similar pressures from habitat loss and direct exploitation in human-modified landscapes (Costa-Pereira & Galetti, 2015). Although birds cannot provide a complete picture of seed dispersal on land cover gradients, they appear to play a major role, especially for later successional species (de la Peña-Domene et al. 2014).

We have argued that patterns of species occurrence across land cover gradients can provide insights into the likely transfer of seeds between habitat types (Sekercioglu, 2006). This is based on the reasoning that the locations in our study represent adjacent portions of a gradient and previous evidence suggests that individual birds will move between adjacent land cover types (Mayhew et al. 2019) with widespread spillover effects (Hatfield et al. 2020). However, this type of information is sparse and species may vary in how often they move between land cover classes. Our approach also makes assumptions about the food plants of frugivorous birds, despite relatively limited knowledge of diets and trophic interactions, particularly in the context of tropical forest birds.

To address these issues, further work is required to clarify rates of movement between land covers by tracking individual birds, and to provide more detailed information about avian diets through techniques such as metabarcoding (Hoenig et al., 2022). This improvement in dietary information would allow for a more comprehensive treatment of species, including abundant generalists, for which fruit is not the main component of their diet but which may nonetheless function as important seed dispersers for particular plant species (Pizo, 2004). Nonetheless, the addition of these species is unlikely to alter our main conclusions because generalist species tend to be more important in degraded forests and are unlikely to provide dispersal services for larger seeds.

Along with more detail on movements and diet, we also require robust data on abundance. Without a better grasp on the number of individual birds involved, it is challenging to evaluate the implications of presence-absence data, as functions are often effectively lost even when particular frugivore species are still present but reduced to such low numbers that they are essentially not delivering any effective seed dispersal function. Future analyses should also encompass the role of seed-dispersing mammals such as primates and bats, which may play a vital role in mediating regeneration potential of human-modified tropical forests (Stoner et al., 2007).

### 4.2 Pathways to resilience in tropical forests

With tropical deforestation continuing at a rapid pace, maintaining the potential for forest regeneration is a critical concern. Logged and disturbed rainforests contain tree species with lower average wood density and smaller seeds (Bello et al., 2015; Hawes et al., 2020) while cleared areas have greatly reduced value for biodiversity. The relevance of strategies promoting rapid and affordable forest regeneration is brought into sharp focus by policy targets, including the Brazilian government’s Nationally Determined Contributions under the Paris Agreement, which represent a binding commitment to achieve carbon neutrality and mitigate climate change impacts, partly through forest restoration. For example, the state of Pará has committed to restore c.5 Mha of Amazonian rainforest by 2030 (Plano Estadual Amazônia Agora, 2020). Such targets can be achieved through direct interventions such as tree-planting programmes. However, the passive restoration of tropical forests via natural processes has a number of benefits over active restoration, including increased diversity (Chazdon & Guariguata, 2016) and greater potential to track climate change (Fricke et al., 2022). In addition, natural regeneration is the only financially viable solution over extensive degraded areas because it can be delivered at minimal cost (Chazdon & Guariguata, 2016).

Where forest restoration is a key objective, our results suggest that dispersal from natural seed sources would require close proximity to intact native forest, supporting earlier work (e.g., Cardaso da Silva et al., 1996). Cleared areas further away from native forests may require alternative approaches, such as applied nucleation and nurse trees, to help encourage seed dispersers to use more of the landscape (Pizo & dos Santos, 2011; Freeman et al., 2021). The likelihood of recruitment will often be low, however, as many matrix environments provide unsuitable terrain for seed germination and sapling growth (Reid & Holl, 2013). Our results also suggest that degraded forest plays a critical role in supplying seeds and seed-dispersal agents. These disturbed habitats not only hold more biodiversity than fully cleared landscapes, they also support landscape-level resilience by providing a vital source and conduit for seed transfer into the non-forest matrix. Thus, prioritising the conservation and management of substantial areas of logged and secondary forests is not only beneficial in terms of their efficient nutrient cycling and carbon uptake (Malhi et al. 2022), but also vital for promoting the healthy seed-dispersal dynamics required for rapid and economical reforestation programmes.

While secondary and disturbed forests can play a key role, our findings highlight the importance of preserving extensive intact forests to promote seed transfer from large-seeded trees, including commercially important hardwoods and other species with tall stature crucial to achieving the vegetation structure and microhabitat complexity of old growth forests (Hawes et al., 2020). In effect, the potential for forest resilience and regeneration will best be delivered by mosaic landscapes with degraded-forest buffers around core areas of intact forest that are not too distant. This tallies with proposals to focus restoration efforts on landscapes with intermediate forest cover levels and good connectivity with intact forest (Tambosi et al., 2014; Mayhew et al., 2019).

## Acknowledgments

This research was funded by Natural Environment Research Council, UK (standard grant NE/K016431/1). JHH was supported by the Leverhulme Centre for Anthropocene Biodiversity (RC-2018-021). We also thank the field teams and landowners who contributed to the generation of the bird survey data. This paper is #XXX in the Rede Amazônia Sustentável publication series.

## References

Ali, J. R., Blonder, B. W., Pigot, A. L., & Tobias, J. A. (2023). Bird extinctions threaten to cause disproportionate reductions of functional diversity and uniqueness. Functional Ecology, 37, 162–175. 10.1111/1365-2435.14201

Athiê, S., & Dias, M. M. (2016). Use of perches and seed dispersal by birds in an abandoned pasture in the Porto Ferreira state park, southeastern Brazil. Brazilian Journal of Biology, 76, 80–92.

Banks-Leite, C., Pardini, R., Boscolo, D., Cassano, C. R., Püttker, T., Barros, C. S., & Barlow, J. (2014). Assessing the utility of statistical adjustments for imperfect detection in tropical conservation science. Journal of Applied Ecology, 51, 849–859. 10.1111/1365-2664.12272

Baselga, A., Orme, D., Villeger, S., De Bortoli, J., Leprieur, F., & Logez, M. (2021). betapart: Partitioning beta-diversity into turnover and nestedness components. R package version 1.5.3. https://cran.r-project.org/package=betapart

Bello, C., Galetti, M., Pizo, M. A., Magnago, L. F. S., Rocha, M. F., Lima, R. A. F., Peres, C. A., Ovaskainen, O., & Jordano, P. (2015). Defaunation affects carbon storage in tropical forests. Science Advances, 1, e1501105. 10.1126/sciadv.1501105

Beltrán, L. C., & Howe, H. F. (2020). The frailty of tropical restoration plantings. Restoration Ecology, 28, 16–21. 10.1111/rec.13066

Biggs, C. R., Yeager, L. A., Bolser, D. G., Bonsell, C., Dichiera, A. M., Hou, Z., Keyser, S. R., Khursigara, A. J., Lu, K., Muth, A. F., Negrete Jr., B., & Erisman, B. E. (2020). Does functional redundancy affect ecological stability and resilience? A review and meta-analysis. Ecosphere, 11, e03184. 10.1002/ecs2.3184

BirdLife International. (2022). IUCN Red List for birds. www.birdlife.org

BirdLife International and Handbook of the Birds of the World. (2020). Bird species distribution maps of the world. Version 2020.1. Available at http://datazone.birdlife.org/species/requestdis.

Bovo, A. A. A., Ferraz, K. M. P. M. B., Magioli, M., Alexandrino, E. R., Hasui, É., Ribeiro, M. C., & Tobias, J. A. (2018). Habitat fragmentation narrows the distribution of avian functional traits associated with seed dispersal in tropical forest. Perspectives in Ecology and Conservation, 16, 90–96. 10.1016/J.PECON.2018.03.004

Bregman, T. P., Lees, A. C., MacGregor, H. E. A., Darski, B., de Moura, N. G., Aleixo, A., Barlow, J., & Tobias, J. A. (2016). Using avian functional traits to assess the impact of land-cover change on ecosystem processes linked to resilience in tropical forests. Proceedings of the Royal Society B: Biological Sciences, 283, 20161289. 10.1098/rspb.2016.1289

Bregman, T. P., Sekercioglu, C. H., & Tobias, J. A. (2014). Global patterns and predictors of bird species responses to forest fragmentation: Implications for ecosystem function and conservation. Biological Conservation, 169, 372–383. 10.1016/j.biocon.2013.11.024

Brooks, T., Tobias, J. A., & Balmford, A. (1999). Deforestation and bird extinctions in the Atlantic Forest. Animal Conservation, 2, 211–222.

Buchanan, G. M., Donald, P. F., & Butchart, S. H. M. (2011). Identifying priority areas for conservation: A global assessment for forest-dependent birds. PLOS ONE, 6, 1–10. 10.1371/journal.pone.0029080

Burns, K. C. (2013). What causes size coupling in fruit–frugivore interaction webs? Ecology, 94, 295–300. 10.1890/12-1161.1

Cadotte, M. W., Carscadden, K., & Mirotchnick, N. (2011). Beyond species: functional diversity and the maintenance of ecological processes and services. Journal of Applied Ecology, 48, 1079–1087. 10.1111/j.1365-2664.2011.02048.x

Camargo, P. H. S. A., Pizo, M. A., Brancalion, P. H. S., & Carlo, T. A. (2020). Fruit traits of pioneer trees structure seed dispersal across distances on tropical deforested landscapes: Implications for restoration. Journal of Applied Ecology, 57, 2329–2339. 10.1111/1365-2664.13697

Canale, G. R., Peres, C. A., Guidorizzi, C. E., Gatto, C. A. F., & Kierulff, M. C. M. (2012). Pervasive defaunation of forest remnants in a tropical biodiversity hotspot. PLOS ONE, 7, e41671. 10.1371/journal.pone.0041671

Cardaso Da Silva, J. M., Uhl, C. & Murray, G. (1996). Plant succession, landscape management, and the ecology of frugivorous birds in abandoned Amazonian pastures. Conservation Biology, 10, 491–503. 10.1046/j.1523-1739.1996.10020491.x

Carlo, T. A., & Morales, J. M. (2016). Generalist birds promote tropical forest regeneration and increase plant diversity via rare-biased seed dispersal. Ecology, 97, 1819–1831. 10.1890/15-2147.1

Chazdon, R. L., & Guariguata, M. R. (2016). Natural regeneration as a tool for large-scale forest restoration in the tropics: prospects and challenges. Biotropica, 48, 716–730. 10.1111/btp.12381

Chazdon, R. L. & Uriarte, M. (2016). Natural regeneration in the context of large-scale forest and landscape restoration in the tropics. Biotropica, 48, 709–715.

Costa-Pereira, R., & Galetti, M. (2015). Frugivore downsizing and the collapse of seed dispersal by fish. Biological Conservation, 191, 809–811.

da Silva, J. M. C., & Tabarelli, M. (2000). Tree species impoverishment and the future flora of the Atlantic forest of northeast Brazil. Nature, 404, 72–74. 10.1038/35003563

de Assis Bomfim, J., Guimarães, P. R., Peres, C. A., Carvalho, G., & Cazetta, E. (2018). Local extinctions of obligate frugivores and patch size reduction disrupt the structure of seed dispersal networks. Ecography, 41, 1899–1909. 10.1111/ecog.03592

de la Peña-Domene, M., Martínez-Garza, C., Palmas-Pérez, S., Rivas-Alonso, E., & Howe, H. F. (2014). Roles of birds and bats in early tropical-forest restoration. PLOS ONE, 9, 1–6. 10.1371/journal.pone.0104656

Dehling, D. M., Töpfer, T., Schaefer, H. M., Jordano, P., Böhning-Gaese, K., & Schleuning, M. (2014). Functional relationships beyond species richness patterns: trait matching in plant–bird mutualisms across scales. Global Ecology and Biogeography, 23, 1085–1093. 10.1111/geb.12193

Dirzo, R., Young, H. S., Galetti, M., Ceballos, G., Isaac, N. J. B., & Collen, B. (2014). Defaunation in the Anthropocene. Science, 345, 401–406. 10.1126/science.1251817

Donatti, C. I., Guimarães, P. R., Galetti, M., Pizo, M. A., Marquitti, F. M. D., & Dirzo, R. (2011). Analysis of a hyper-diverse seed dispersal network: modularity and underlying mechanisms. Ecology Letters, 14, 773–781. 10.1111/j.1461-0248.2011.01639.x

Freeman, A. N. D., Freebody, K., Montenero, M., Moran, C., Shoo, L. P., & Catterall, C. P. (2021). Enhancing bird-mediated seed dispersal to increase rainforest regeneration in disused pasture – A restoration experiment. Forest Ecology and Management, 479, 118536. 10.1016/j.foreco.2020.118536

Fricke, E. C., Ordonez, A., Rogers, H. S., & Svenning, J.-C. (2022). The effects of defaunation on plants’ capacity to track climate change. Science, 375, 210–214.

Galetti, M., Guevara, R., Cortes, M. C., Fadini, R., Von Matter, S., Leite, A. B., Labecca, F., Ribeiro, T., Carvalho, C. S., Collevatti, R. G., Pires, M. M., Guimaraes, P. R., Brancalion, P. H., Ribeiro, M. C., & Jordano, P. (2013). Functional extinction of birds drives rapid evolutionary changes in seed size. Science, 340, 1086–1090.

Gardner, C. J., Bicknell, J. E., Baldwin-Cantello, W., Struebig, M. J., & Davies, Z. G. (2019). Quantifying the impacts of defaunation on natural forest regeneration in a global meta-analysis. Nature Communications, 10, 4590. 10.1038/s41467-019-12539-1

Gardner, T. A., Ferreira, J., Barlow, J., Lees, A. C., Parry, L., Vieira, I. C. G., Berenguer, E., Abramovay, R., Aleixo, A., Andretti, C., Aragão, L. E. O. C., Araújo, I., de Ávila, W. S., Bardgett, R. D., Batistella, M., Begotti, R. A., Beldini, T., de Blas, D. E., Braga, R. F.,…Zuanon, J. (2013). A social and ecological assessment of tropical land uses at multiple scales: the Sustainable Amazon Network. Philosophical Transactions of the Royal Society of London. Series B, Biological Sciences, 368, 20120166. 10.1098/rstb.2012.0166

González-Varo, J. P., Carvalho, C. S., Arroyo, J. M., & Jordano, P. (2017). Unravelling seed dispersal through fragmented landscapes: Frugivore species operate unevenly as mobile links. Molecular Ecology, 26, 4309–4321. 10.1111/mec.14181

Habel, J. C., Tobias, J. A., & Fischer, C. (2019). Movement ecology of Afrotropical birds: Functional traits provide complementary insights to species identity. Biotropica, 51, 894–902. 10.1111/btp.12702

Hatfield, Jack H., Barlow, J., Joly, C. A., Lees, A. C., Parruco, C. H. de F., Tobias, J. A., Orme, C. D. L., & Banks-Leite, C. (2020). Mediation of area and edge effects in forest fragments by adjacent land use. Conservation Biology, 34, 395–404. 10.1111/cobi.13390

Hawes, J. E., Vieira, I. C. G., Magnago, L. F. S., Berenguer, E., Ferreira, J., Aragão, L. E. O. C., Cardoso, A., Lees, A. C., Lennox, G. D., Tobias, J. A., Waldron, A., & Barlow, J. (2020). A large-scale assessment of plant dispersal mode and seed traits across human-modified Amazonian forests. Journal of Ecology, 108, 1373–1385. 10.1111/1365-2745.13358

Heleno, R. H., Ross, G., Everard, A., Memmott, J., & Ramos, J. A. (2011). The role of avian ‘seed predators’ as seed dispersers. Ibis, 153, 199–203. 10.1111/j.1474-919X.2010.01088.x

Hoenig, B. D., Snider, A. M., Forsman, A. M., Hobson, K. A., Latta, S. C., Miller, E. T., Polito, M. J., Powell, L. L., Rogers, S. L., Sherry, T. W., Toews, D. P. L., Welch, A. J., Taylor, S. S., & Porter, B. A. (2022). Current methods and future directions in avian diet analysis. Ornithology, 139, 1–28. 10.1093/ornithology/ukab077

Hurlbert, A. H., & Jetz, W. (2007). Species richness, hotspots, and the scale dependence of range maps in ecology and conservation. Proceedings of the National Academy of Sciences, 104, 13384–13389. 10.1073/pnas.0704469104

Jetz, W., Sekercioglu, C. H., & Watson, J. E. M. (2008). Ecological Correlates and Conservation Implications of Overestimating Species Geographic Ranges. Conservation Biology, 22, 110–119. 10.1111/j.1523-1739.2007.00847.x

Joly, C. A., Metzger, J. P., & Tabarelli, M. (2014). Experiences from the Brazilian Atlantic Forest: ecological findings and conservation initiatives. New Phytologist, 204, 459–473.

Lees, A. C., Aleixo, A., Paraense, M., Goeldi-Mpeg, E., Barlow, J., & Berenguer, E. (2012). Paragominas: A quantitative baseline inventory of an eastern Amazonian avifauna. Revista Brasileira De Ornitologia, 20, 93–118.

Lees, A. C., Moura, N. G. de, Andretti, C. B., Davis, B. J. W., Lopes, E. V., Henriques, L. M. P., Aleixo, A., Barlow, J., Ferreira, J., & Gardner, T. A. (2013). One hundred and thirty-five years of avifaunal surveys around Santarém, central Brazilian Amazon. Revista Brasileira De Ornitologia, 21, 16–57.

Lees, A. C., & Pimm, S. L. (2015). Species extinct before we know them? Current Biology, 25, R177–R180.

Malhi, Y., Riutta, T., Wearn, O. R., Deere, N. J., Mitchell, S. L., Bernard, H., Majalap, N., Nilus, R., Davies, Z. G., Ewers, R. M., & Struebig, M. J. (2022). Logged tropical forests have amplified and diverse ecosystem energetics. Nature, 612, 707–713. 10.1038/s41586-022-05523-1

Mason, N. W. H., Mouillot, D., Lee, W. G., & Wilson, J. B. (2005). Functional richness, functional evenness and functional divergence: the primary components of functional diversity. Oikos, 111, 112–118. 10.1111/j.0030-1299.2005.13886.x

Mayhew, R. J., Tobias, J. A., Bunnefeld, L., & Dent, D. H. (2019). Connectivity with primary forest determines the value of secondary tropical forests for bird conservation. Biotropica, 51, 219–233. 10.1111/btp.12629

McFadden, I. R., Fritz, S. A., Zimmermann, N. E., Pellissier, L., Kissling, W. D., Tobias, J. A., Schleuning, M., & Graham, C. H. (2022). Global plant-frugivore trait matching is shaped by climate and biogeographic history. Ecology Letters, 25, 686–696. 10.1111/ele.13890

Moura, N. G., Lees, A. C., Aleixo, A., Barlow, J., Berenguer, E., Ferreira, J., Mac Nally, R., Thomson, J. R., & Gardner, T. A. (2016). Idiosyncratic responses of Amazonian birds to primary forest disturbance. Oecologia, 180, 903–916. 10.1007/s00442-015-3495-z

Moura, N. G., Lees, A. C., Andretti, C. B., Davis, B. J. W., Solar, R. R. C., Aleixo, A., Barlow, J., Ferreira, J., & Gardner, T. A. (2013). Avian biodiversity in multiple-use landscapes of the Brazilian Amazon. Biological Conservation, 167, 339–348. 10.1016/J.BIOCON.2013.08.023

Nunes, C. A., Berenguer, E., França, F., Ferreira, J., Lees, A. C., Louzada, J., Sayer, E. J., Solar, R., Smith, C. C., Aragão, L. E. & Braga, D. D. L. (2022). Linking land-use and land-cover transitions to their ecological impact in the Amazon. Proceedings of the National Academy of Sciences, 119, e2202310119. 10.1073/pnas.22023101

Orme, C. D. L., Mayor, S., dos Anjos, L., Develey, P. F., Hatfield, J. H., Morante-Filho, J. C., Tylianakis, J. M., Uezu, A., & Banks-Leite, C. (2019). Distance to range edge determines sensitivity to deforestation. Nature Ecology and Evolution, 3, 886–891. 10.1038/s41559-019-0889-z

Plano Estadual Amazônia Agora. (2020). Secretaria de Estado de Meio Ambiente e Sustentabilidade - Semas, DECRETO N° 941, DE 03 DE AGOSTO DE 2020. https://www.semas.pa.gov.br/legislacao/publico/view/8457

Pérez-Méndez, N., Jordano, P., García, C., & Valido, A. (2016). The signatures of Anthropocene defaunation: cascading effects of the seed dispersal collapse. Scientific Reports, 6, 24820. 10.1038/srep24820

Petchey, O. L., & Gaston, K. J. (2002). Functional diversity (FD), species richness and community composition. Ecology Letters, 5, 402–411.

Pigot, A. L., Bregman, T., Sheard, C., Daly, B., Etienne, R. S., & Tobias, J. A. (2016). Quantifying species contributions to ecosystem processes: A global assessment of functional trait and phylogenetic metrics across avian seed-dispersal networks. Proceedings of the Royal Society B: Biological Sciences, 283, 20161597. 10.1098/rspb.2016.1597

Pigot, A. L., Sheard, C., Miller, E. T., Bregman, T. P., Freeman, B. G., Roll, U., Seddon, N., Trisos, C. H., Weeks, B. C., & Tobias, J. A. (2020). Macroevolutionary convergence connects morphological form to ecological function in birds. Nature Ecology & Evolution, 4, 230–239.

Pizo, M.A. (2004). Frugivory and habitat use by fruit-eating birds in a fragmented landscape of southeast Brazil. Ornitologia neotropical, 15, 117–126.

Pizo, M. A., & dos Santos, B. T. P. (2011). Frugivory, post-feeding flights of frugivorous birds and the movement of seeds in a Brazilian fragmented landscape. Biotropica, 43, 335–342. 10.1111/j.1744-7429.2010.00695.x

Rehm, E. M., Chojnacki, J., Rogers, H. S., & Savidge, J. A. (2017). Differences among avian frugivores in seed dispersal to degraded habitats. Restoration Ecology, 26, 760–766. 10.1111/rec.12623

Reid, J. L., & Holl, K. D. (2013). Arrival ≠ Survival. Restoration Ecology, 21, 153–155. 10.1111/j.1526-100X.2012.00922.x

Reid, J. L., Zahawi, R. A., Zárrate-Chary, D. A., Rosales, J. A., Holl, K. D., & Kormann, U. (2021). Multi-scale habitat selection of key frugivores predicts large-seeded tree recruitment in tropical forest restoration. Ecosphere, 12, e03868. 10.1002/ecs2.3868

Robinson, W. D., Lees, A. C., & Blake, J. G. (2018). Surveying tropical birds is much harder than you think: a primer of best practices. Biotropica, 50, 846–849. 10.1111/btp.12608

Schleuning, M., Neuschulz, E. L., Albrecht, J., Bender, I. M. A., Bowler, D. E., Dehling, D. M., Fritz, S. A., Hof, C., Mueller, T., Nowak, L., Sorensen, M. C., Böhning-Gaese, K., & Kissling, W. D. (2020). Trait-based assessments of climate-change impacts on interacting species. Trends in Ecology & Evolution, 35, 319–328. 10.1016/j.tree.2019.12.010

Sekercioglu, C. H. (2006). Increasing awareness of avian ecological function. Trends in Ecology & Evolution, 21, 464–471. 10.1016/j.tree.2006.05.007

Sethi, P. I. A., & Howe, H. F. (2009). Recruitment of hornbill-dispersed trees in hunted and logged forests of the Indian eastern Himalaya. Conservation Biology, 23, 710–718. 10.1111/j.1523-1739.2008.01155.x

Sheard, C., Neate-Clegg, M. H. C., Alioravainen, N., Jones, S. E. I., Vincent, C., MacGregor, H. E. A., Bregman, T. P., Claramunt, S., & Tobias, J. A. (2020) Ecological drivers of global gradients in avian dispersal inferred from wing morphology. Nature Communications, 11, 2463.

Solar, R. R. C., Barlow, J., Ferreira, J., & Berenguer, E. (2015). How pervasive is biotic homogenization in human modified tropical forest landscapes? Ecology, 18, 1108–1118. 10.1111/ele.12494

Stoner, K.E., Riba-Hernández, P., Vulinec, K. & Lambert, J.E. (2007). The role of mammals in creating and modifying seedshadows in tropical forests and some possible consequences of their elimination. Biotropica, 39, 316–327.

Tambosi, L. R., Martensen, A. C., Ribeiro, M. C., & Metzger, J. P. (2014). A framework to optimize biodiversity restoration efforts based on habitat amount and landscape connectivity. Restoration Ecology, 22, 169–177.

Tella, J.L., Baños-Villalba, A., Hernández-Brito, D., Rojas, A., Pacífico, E., Díaz-Luque, J.A., Carrete, M., Blanco, G., & Hiraldo, F. (2015). Parrots as overlooked seed dispersers. Frontiers in Ecology and the Environment, 13, 338–339. 10.1890/1540-9295-13.6.338

Tobias, J.A., Ottenburghs, J., & Pigot, A. (2020). Avian diversity: speciation, macroevolution and ecological function. Annual Reviews of Ecology, Evolution and Systematics, 51, 533–560.

Tobias, J. A., Sheard, C., Pigot, A. L., Devenish, A. J. M., Yang, J., Sayol, F., Neate-Clegg, M. H. C., Alioravainen, N., Weeks, T. L., Barber, R. A., Walkden, P. A., MacGregor, H. E. A., Jones, S. E. I., Vincent, C., Phillips, A. G., Marples, N. M., Montaño-Centellas, F. A., Leandro-Silva, V., Claramunt, S.,…Schleuning, M. (2022). AVONET: morphological, ecological and geographical data for all birds. Ecology Letters, 25, 581–597. 10.1111/ele.13898

Trisos, C. H., Petchey, O. L., & Tobias, J. A. (2014). Unraveling the interplay of community assembly processes acting on multiple niche axes across spatial scales. The American Naturalist, 184, 593–608. 10.1086/678233

Villéger, S., Mason, N. W. H., & Mouillot, D. (2008). New multidimensional functional diversity indices for a multifaceted framework in functional ecology. Ecology, 89, 2290–2301. 10.1890/07-1206.1

Wheelwright, N. T. (1985). Fruit-size, gape width, and the diets of fruit-eating birds. Ecology, 66, 808–818. 10.2307/1940542

Wilman, H., Belmaker, J., Simpson, J., de la Rosa, C., Rivadeneira, M. M., & Jetz, W. (2014). EltonTraits 1.0: Species-level foraging attributes of the world’s birds and mammals. Ecology, 95, 2027. 10.1890/13-1917.1

